# Plant–soil feedback persists beyond host death to shape density-dependent plant competition

**DOI:** 10.1101/2025.10.04.680482

**Authors:** Ching-Lin Huang, Joe Wan, Shuo Wei, Chia-Hao Chang-Yang, Po-Ju Ke

**Affiliations:** Institute of Ecology and Evolutionary Biology, National Taiwan University, Taipei, Taiwan; Department of Ecology, Evolution and Behavior, University of Minnesota-Twin Cities, St. Paul, Minnesota, USA; Yale School of the Environment, Yale University, New Haven, Connecticut, USA; Department of Biological Sciences, National Sun Yat-sen University, Kaohsiung, Taiwan

**Author notes:** These authors contributed equally (Ching-Lin Huang, Joe Wan).

**Keywords:** Forest dynamic plot, Invasion growth rate, Legacy effect, Mycorrhizal fungi, Nonlinear density dependence, Subtropical forest

## Abstract

**Background and Aims:** Plant–soil feedback (PSF) shapes plant competition, yet classic PSF experiments often overlook the density dependence and temporal complexities of PSFs, especially after host death. We ask whether microbial effects vary after host death and mediate density dependence in seedling competition.

**Methods:** We combined forest inventory data from a subtropical forest with a density-gradient greenhouse experiment to evaluate seedling competition among two tree species with different mycorrhizal associations (ectomycorrhizal versus arbuscular mycorrhizal). In 2023, we collected soil inocula from living and dead trees (died between 2014 and 2019) with unsterilized and sterilized treatments. We applied invasion analysis to infer seedling competitive outcomes and used bootstrapping to evaluate uncertainty.

**Results:** Soil microbial communities shaped seedling competition, favoring the ectomycorrhizal species. Using soils collected from living versus dead hosts as live inocula, we found similar plant competitive outcomes, indicating that PSF persists for over 4–9 years following host death. In contrast, when using sterilized inoculum, we found shifts in competition after host death, suggesting underlying abiotic changes which might masked by microbial effects. Moreover, we found significant evidence that soil microbes can mediate nonlinear density dependence in seedling competition.

**Conclusion:** We provide experimental evidence of persistent microbial legacies that benefit ectomycorrhizal tree species in a subtropical forest. Our study demonstrates how integrating field-based census data with density-gradient experiments and explicit uncertainty estimation can better capture the temporal dimensions, complexities of density dependence, and uncertainties of PSF.

## Introduction

Reciprocal interactions between plants and soil microbes, known as plant–soil feedback (PSF), shape the structure and dynamics of plant communities (Bever et al., 1997, 2010, van der Putten et al., 2013). Plants, as hosts, condition the soil microbial community differently, altering the relative abundance of beneficial and detrimental soil microbes among host plants (Bever et al., 2012, Hu et al., 2018). In turn, these shifts in plant species-specific soil microbial composition influence the growth of nearby plant individuals and the establishment of new recruits (Bever et al., 1997, Bever, 2003). As plants vary in their responses to soil microbial communities, PSF can alter competitive dynamics by modifying the relative competitive abilities among plants (Ke and Wan, 2020, Kandlikar et al., 2019), thereby shaping their relative abundance patterns (Klironomos, 2002, Mangan et al., 2010) and assembly dynamics (Fukami and Nakajima, 2013). Despite the widely recognized importance of PSF for plant community dynamics, the density dependence and temporal complexities of PSF remain underexplored (Gundale and Kardol, 2021, Chung, 2023, Ke et al., 2024). These complexities are expected since soil microbes respond to host density and turn over through time (Steinauer et al., 2023, Willing et al., 2024). Resolving the density- and time-dependencies of PSF is essential for explaining and predicting plant coexistence and community dynamics.

To understand how soil microbes influence plant competitive outcome, studies estimate theory-derived indices by growing plants in soils conditioned by different plant species (Bever et al., 1997, Crawford et al., 2019, Yan et al., 2022). However, these experiments typically measure the responses of individual plants in isolation, implicitly assuming that soil microbes only modify the intrinsic growth rate of plant populations (Bever et al., 1997, Kandlikar, 2024). This classical assumption has been challenged by experiments showing that microbial effects can vary with plant density, indicating density dependence in plant–soil microbe interactions (West, 1996, Willing et al., 2024). Moreover, this approach overlooks the long-standing recognition that plant–plant interactions are inherently density-dependent; that is, plant performance is influenced by the abundance of neighboring plants (Watkinson and Freckleton, 1996). While some recent studies have employed various experimental designs to quantify how soil microbes mediate the density-dependence of plant–plant interaction (e.g., Chung and Rudgers, 2016, Cardinaux et al., 2018, Huangfu et al., 2022), we lack practical guidelines to address this aspect of PSF. To this end, Ke and Wan (2023) proposed a theory-grounded density-gradient experimental design to measure density-dependent microbial effects on plant performance, which can also detect nonlinearity in density dependence. This framework offers a tractable approach for assessing how soil microbes influence plant competitive outcomes by altering the form of density-dependent plant–plant interactions.

Recent studies have also underscored the importance of the temporal complexity of plant– soil microbe interactions (Kardol et al., 2013, Gundale and Kardol, 2021, Chung, 2023, Ke et al., 2024). Specifically, to better predict the impact of conditioned microbial communities on newly arriving seedlings, it is essential to examine how microbial effects develop over a conditioning plant’s lifespan and how they vary after host death. Theoretical studies have shown that persistent microbial legacies are crucial for PSF to maintain plant diversity (Ke et al., 2021, Miller and Allesina, 2021); however, empirical studies have yielded contrasting results regarding the persistence of these microbial legacies. For instance, Bennett et al. (2022) used field soils from living or dead tree hosts to show strong and consistent legacy effects on seedling demography. However, Ou et al. (2024) demonstrated that a six-month temporal lag between conditioning and planting resulted in different predictions of competitive outcomes, suggesting that microbial effects are not constant after host mortality. Moreover, while a recent study using forest inventory data suggests that density-dependent plant–plant interactions are persistent after tree death and implicates a potential role for soil microbes (Magee et al., 2024), experimental studies that directly test the microbial mechanisms underlying such patterns are still lacking.

Here, we explore how density-dependent plant–plant interactions are influenced by soil microbial legacies left by dead host plants, testing the hypothesis that the effects of soil microbes on plant competitive outcome vary through time. To address this, we combine forest inventory data with a density-gradient greenhouse experiment (modified from Ke and Wan, 2023), using soils collected beneath living and dead trees to inoculate a seedling competition experiment. We leverage a well-studied field system: Fushan Forest Dynamic Plot (Fushan FDP), a subtropical forest in Taiwan with four complete forest inventories. Given that preparing soils representing multiple years since host death is often infeasible in greenhouse settings, we use field soil beneath living and dead trees to study the temporal changes of PSF by leveraging different host tree statuses in nature.

Our experiment monitors seedling competition under a density-gradient experiment design to estimate population-level intrinsic growth rates and density-dependent plant–plant interactions, either with or without microbial mediation. Finally, we apply invasion analysis to evaluate species persistence and competitive outcomes, and compare how these outcomes shift across microbial treatments and over time.

## Materials and Methods

### Study species and field soil preparation

We studied a pair of common subtropical low-montane forest trees, *Engelhardia roxburghiana* (Juglandaceae) and *Machilus zuihoensis* (Lauraceae). Both are among the most abundant species at Fushan Forest Dynamics Plot (Fushan FDP; 24°45’40” N, 121°33’28” E; elevation 600–733 m a.s.l.). Namely, *E. roxburghiana* is the most frequent canopy tree species by number of individuals (270.6 individuals/ha; 17.0% of all canopy-tree individuals) and *M. zuihoensis* is the 7th most frequent (100.2 individuals/ha; 6.3% of all canopy-tree individuals). In terms of above-ground biomass, *M. zuihoensis* ranks 4th (15.0 Mg/ha), while *E. roxburghiana* ranks 8th (6.2 Mg/ha). From 2004 to 2024, the annual mortality rate was approximately 3.6% for *E. roxburghiana* and 2.1% for *M. zuihoensis*, mainly due to crown damage (Zuleta et al., 2022). While both study species are large canopy trees, they are likely to interact differently with soil microbes. For instance, *E. roxburghiana* is capable of forming ectomycorrhizae (Haug et al., 1994), while *M. zuihoensis* associates only with arbuscular mycorrhizal fungi. Moreover, they differ in their phylogenetic relatedness to surrounding vegetation: *E. roxburghiana* is the only member of the Juglandaceae at the site, whereas *M. roxburghiana* is one of nearly a dozen species in the Lauraceae at Fushan FDP (including 4 out of the 10 most common canopy trees).

Field soils were collected from the Fushan FDP in late November 2023. For both species, we randomly selected 10 individuals in each of two statuses: living (individuals that were alive at the time of sampling) and dead (individuals that had died between 2014 and 2019, i.e., had been dead for 4 to 9 years, according to forest census data; Fig. S1). To ensure that dead individuals could be accurately located in the field and control for tree age, we restricted our sampling to relatively mature individuals in the 2019 tree census (diameter at breast height 14.5–38 cm; average: 24.5 cm). Note that only nine living *M. zuihoensis* individuals were sampled, as one individual was found dead during sampling. Using a surface-sterilized soil core sampler (0.5% bleach followed by 75% ethanol), we collected 1.0 L of topsoil (∼10 cm depth, excluding litter; multiple soil sampling points per tree) from within a 1 m radius of each selected individual. Soils from different individuals were kept in separate sealed sterile Ziploc bags, transported to the lab in a cooler, sieved (2 mm mesh) to remove stones and root fragments, and stored at 4 °C before being used in the greenhouse experiment.

A mixed soil sampling approach (Gundale et al., 2017) was used to prepare the soil inoculum for our greenhouse experiment. Specifically, we mixed and homogenized equal volumes of soil collected from all individuals of the same species and status (living or dead). This soil mixture was then mixed with autoclaved perlite following a 6:1 soil:perlite volume ratio to improve drainage, forming the inocula used in our greenhouse experiment. As a baseline for assessing the potential effects of soil microbes, we sterilized half of the mixed soil inocula by autoclaving (121°C and 1.2 kg/cm^2^ for 30 min, incubation at room temperature for 24 h, and another 121°C and 1.2 kg/cm^2^ for 30 min). Overall, for each species, our experiment included four different inoculum types, defined by the two conditioning host plant statuses (living or dead based on forest census) and the two sterilization treatments (unsterilized or sterilized).

### Density-gradient greenhouse experiment

Our greenhouse experiment was conducted according to the following procedure. We germinated seedlings of *M. zuihoensis* and *E. roxburghiana* in growth chambers. Seeds of both species were purchased from a seed supplier, surface-sterilized with 0.5% bleach solution, and rinsed with reverse osmosis water before germination. Since the seeds of *M. zuihoensis* experience physiological dormancy (Chien et al., 1994), the seeds were cold stratified at 4 °C for two months in sealed sterile Ziploc bags filled with sterilized sphagnum moss. In October 2023, once the *M. zuihoensis* seeds started to germinate, seedlings with extended primary leaves were transferred to trays (8 cm in depth) filled with sterilized perlite to avoid root girdling. Meanwhile, we directly sowed *E. roxburghiana* seeds on identical trays as *M. zuihoensis*. Both species were placed in growth chambers maintained at 25/20 °C day/night temperature, 12/12 hrs day/night photoperiod (light intensity ∼150 *µ*mol · m^−2^ · s^−1^), and a constant 90% relative humidity. Seeds and seedlings were misted daily with reverse osmosis water to maintain substrate moisture.

Following the experimental design proposed in Ke and Wan (2023), we implemented a full factorial design consisting of the two plant focal species × two plant competitors × four competitor density scenarios × two field host plant statuses (living or dead) × two soil sterilization treatments (unsterilized or sterilized), with four replicate pots for each combination (a total of 256 pots, Fig. 1a). In December 2023, similarly sized seedlings of *M. zuihoensis* and *E. roxburghiana* were transplanted into pots (10.6 cm in top diameter, 16.0 cm in height, ∼ 1 L in volume). Each pot contained one focal individual along with a designated number (0, 1, 2, or 4) of conspecific or heterospecific competitors. The pots were filled with 0.8 L of sterilized planting substrate (peat:perlite = 7:1, by volume), then with 0.1 L of the designated soil inoculum, and finally topped with another 0.1 L of sterilized planting substrate. As this experiment is designed to quantify the growth of each species when it is rare (i.e., invasion growth), the inoculum always matched the competitor species. For example, if the competitor was *M. zuihoensis*, we used the inoculum collected from *M. zuihoensis* in the field. The setup thereby simulates the microbial effects conditioned by a common competitor species (i.e., the resident) as part of its impact on the rare focal species (i.e., the invader). To ensure future identification, we marked every focal individual with a 5 cm plastic stirrer. The whole planting process was completed within three days. Immediately after planting, 15 additional seedlings of each species from the germination trays were oven-dried at 70 °C for 72 hours to measure their average initial dry biomass. To minimize transplant shock, we began the experiment in growth chambers under identical settings to those used for seed germination. After 80 days, as plants began to fill the growth chamber space, we transferred all pots to the phytotron at National Taiwan University under 25/20 °C day/night temperature and a natural photoperiod. Throughout the experiment, plants were watered with 30–50 mL of reverse osmosis water two to three times per week, and were regularly weeded to minimize the effects of non-target species on seedling performance.

**Figure 1.**
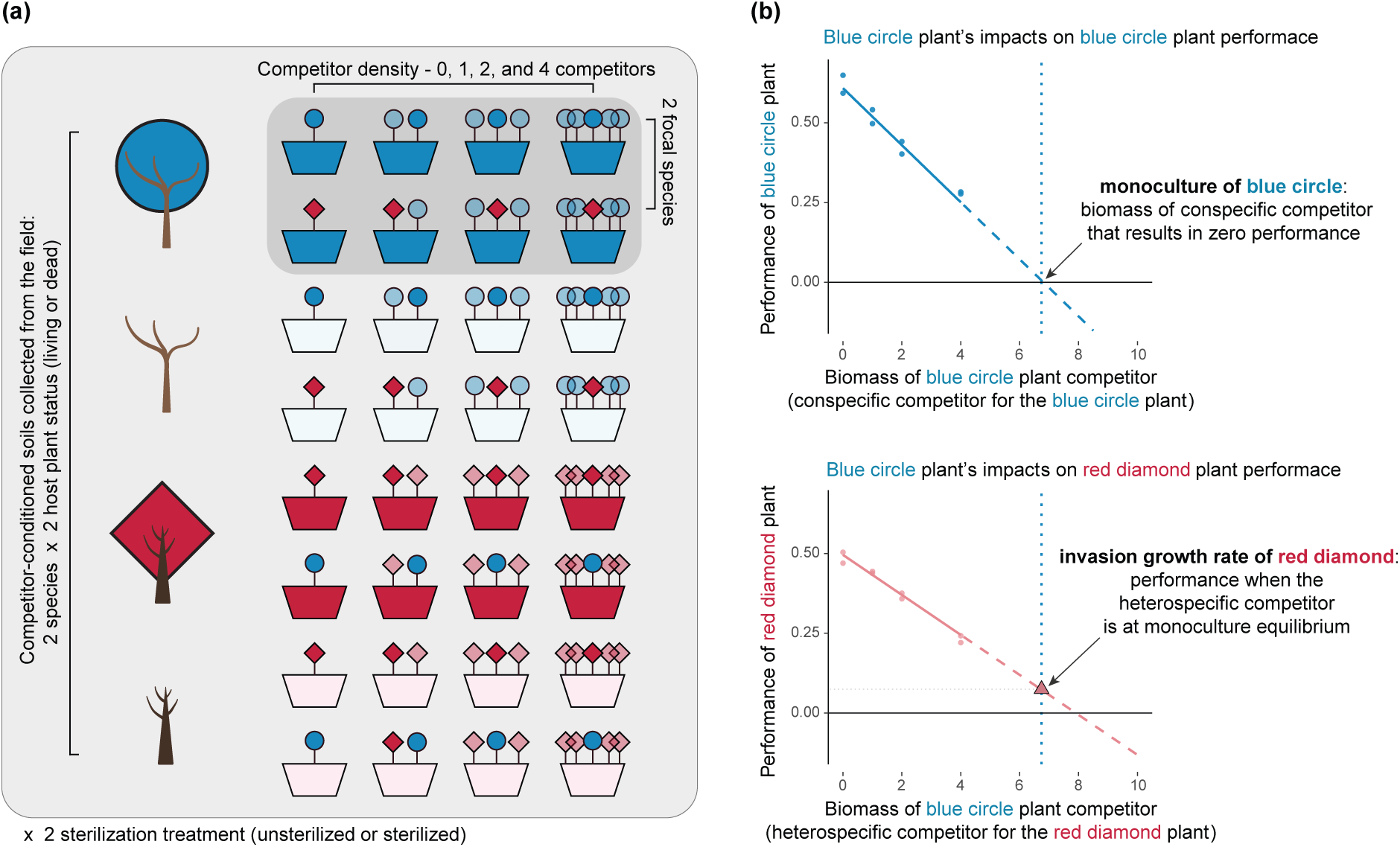
Experimental design and schematic diagram for the invasion analysis following Ke and Wan (2023). (a) The experiment measures the performance (growth) of a focal individual (fully opaque blue circle or red diamond plants, representing the two tree species considered in our study) competing against different densities of competitors (translucent individuals). We added competitor-conditioned soils collected from either living or dead tree individuals in the field (inocula from different tree statuses are represented as pots with opaque and translucent colors, respectively; see also tree icons on the left representing host tree species and status). The depicted setup was repeated for two sterilization treatments (i.e., either unsterilized or sterilized inoculum) and was replicated four times, resulting in a total of 256 pots. (b) An example of how to quantify the invasion growth rate for the red diamond plant, using pots within the highlighted gray box in (a). Both diagrams show a regression model relating focal species growth to competitor biomass, with the dashed portion indicating extrapolated range. First, the impact of the blue circle plant (the competitor, with biomass drawn on the x-axis) on conspecific performance (upper panel) provides an estimation of its monoculture equilibrium (vertical blue dotted lines). Second, the performance of the red diamond plant (lower panel) when the blue plant is at its monoculture equilibrium represents the red diamond plant’s invasion growth rate (red triangle).

After six months, we harvested the above- and below-ground tissues of each individual plant between late June and early July 2024. We gently washed all soil from root systems and separated the above- and below-ground tissues based on the position of the highest secondary root. Entangled fine roots from different individuals were carefully separated. We retained as many root fragments as possible and weighed them separately (see section “Data analysis” for how root fragments contribute to calculating individual biomass). Finally, we oven-dried above- and below-ground tissues at 70 °C for at least 72 hours and summed their weights to determine total dry biomass per focal individual. We observed 21 dead individuals, including twelve competitors and nine focal individuals, out of 704 individuals (overall mortality rate: 3%). The above- and below-ground tissues of these dead individuals were harvested using the same method as the surviving ones.

### Data analysis

To examine how plant competition is affected by soil microbes, we first analyzed how focal species’ growth responded to competitor biomass in different inocula using independent second-order polynomial regressions. In these regressions, the intercept term estimates the focal species’ growth in the absence of competitors, whereas the linear and quadratic terms estimate the total density-dependent impact of competitors on the focal species’ growth. Second, based on these regressions, we assessed species persistence and coexistence under different inoculum types using invasion analysis. Third, we evaluated the uncertainty in model parameters and competitive outcomes by bootstrapping. Finally, based on bootstrap distributions, we compared the differences in competitive outcomes between unsterilized versus sterilized inocula and soils from living versus dead hosts to infer how microbial effects on plant competition outcomes change over time. Here, because growth and mortality of seedlings varied among pots, we report the effects of competitor biomass on focal species’ growth as a measure of realized competitive pressure. Nevertheless, results were qualitatively similar when we repeated all analyses using competitor density as the predictor (see Appendix). Detailed methods for each step are as follows.

First, we analyzed how each focal species’ growth responded to competitor biomass in each inoculum type. For each combination of competitor species, field host status (living versus dead), and inoculum sterilization (unsterilized versus sterilized), we fit an independent second-order polynomial regression.

We defined the growth of a focal individual as the log-ratio of its harvested dry biomass to the species’ average initial dry biomass, serving as a proxy of the species’ per-capita population growth rate. This focal individual growth was regressed against competitor biomass independently for each combination of competitor species, field host status, and inoculum sterilization treatment.

Specifically, we considered a second-order polynomial regression model *y* = *r* + *αx* + *βx*^2^ + *ɛ*, where *y* represents the focal individual growth, and *x* is the log-transformed harvested dry biomass of the competitors (i.e., log(1 + biomass*_harvest_*/biomass*_initial_*)). The intercept *r* represents the growth of the focal species in the absence of competitors. The coefficients *α* and *β* represent the linear and quadratic competitor biomass effects on the focal species’ growth. The error term *ɛ* was assumed to follow a normal distribution. To select better-fitted models, we conducted F-tests to compare across nested models and retained the quadratic term only when it significantly improved model fit (AIC and BIC values shown in the Appendix). We kept the linear terms in all models regardless of their significance. This is because theoretical studies have highlighted the importance of the ratio between conspecific and heterospecific competitive effects for inferring competitive outcomes (Adler et al., 2018, Ke and Wan, 2023). As our main goal was to investigate competitive outcomes across different inocula, rather than to test the competitor effect itself, we avoided dropping the linear competitor effects during model selection (Van Dyke et al., 2024) and instead evaluated their significance using bootstrapping, as described below.

Second, we conducted invasion analyses following Ke and Wan (2023) to determine the competitive outcomes between *M. zuihoensis* and *E. roxburghiana* across inocula differing in host statuses and sterilization treatments. This approach estimates the sign and relative magnitude of each species’ invasion growth rate (IGR), which is the population growth rate of a focal species when its competitor (the resident) has reached its monoculture equilibrium. To obtain the mono-culture equilibrium for each species, we extrapolated the fitted regression model to determine the *conspecific* competitor biomass at which growth of the species equals zero (vertical dotted blue line in Fig. 1b). As a proxy for a focal species’ IGR, we used the fitted model to predict the focal species’ growth when its *heterospecific* competitor was at monoculture equilibrium (red triangle in the lower panel of Fig. 1b). A special case can arise when the linear conspecific competitor effect is positive, and no nonlinear term is retained during model selection. In this case, the species’ growth is predicted to increase without bound as conspecific competitor biomass increases, indicating that the competitor biomass required to generate competitive limitation lies beyond our experimental design and the species’ monoculture equilibrium cannot be determined. When the heterospecific competitor’s monoculture equilibrium could not be determined, we assigned the sign of the focal species’ IGR based on the sign of the highest-order heterospecific effect included in the model. Although the exact magnitude of the IGR cannot be determined in such cases, the sign of the IGR alone is sufficient to infer species persistence and competitive outcomes. Therefore, for each inoculum, we qualitatively inferred species persistence and overall competitive outcome from the pair of IGR signs. A positive IGR indicates that the species can persist under competition, whereas a negative IGR suggests that the species would be outcompeted. If both species have positive IGRs, they are predicted to coexist; if only one species has positive IGR, that species is predicted to exclude its competitor; if both species have negative IGRs, they are inferred to show priority effects, where the outcome depends on the initial abundances of the two species.

Third, we assessed the robustness of the invasion analysis results by non-parametric bootstrapping. This allowed us to account for uncertainty in the estimated competitive outcomes, as recently suggested by Terry and Armitage (2024). Specifically, within each combination of focal species, competitor species, host status, and sterilization treatment, we resampled the 16 data points with replacement, consisting of four competitor densities with four replicates, while maintaining the original sample size. Using the same model structure identified during model selection, we re-estimated model parameters (i.e., the growth without competitors *r*, linear competitor effect *α*, and quadratic competitor effect *β*). We then repeated the invasion analysis for each bootstrap sample and recorded the resulting competitive outcomes. This procedure was repeated 10,000 times to generate the bootstrap distributions for the model parameters and competitive outcomes. The 90% confidence intervals for all model parameters were calculated using the percentile method (Davison and Hinkley, 1997). Significant differences in model parameters across inocula were determined by non-overlapping confidence intervals. Furthermore, we used the bootstrap distributions to evaluate the robustness of the inferred competitive outcomes. A positive IGR was regarded as a robust inference if *>* 95% of bootstrap samples yielded a positive value (i.e., a one-sided test at the *α* = 0.05 significance level), suggesting strong evidence for persistence. Similarly, we regarded a competitive outcome as robust if it received over 95% support across the bootstrap replicates. Finally, to infer microbial effects on seedling competition, we compared competitive outcomes between unsterilized versus sterilized inocula and soils collected from living versus dead hosts. Here, given there is no established convention for significance testing of competitive outcomes (Terry and Armitage, 2024), we follow recent practice (Van Dyke et al., 2024, Willing et al., 2024) by comparing proportions of bootstrap support for each competitive outcome to convey the robustness of our inferences.

## Results

### Significance and uncertainties of model parameters

Our model selection process identified a significantly negative quadratic term only in the case when *E. roxburghiana* (focal species) competed with *M. zuihoensis* (competitor) in sterilized inoculum collected under dead hosts (*F* = 8.279, *P* = 0.013; Table S1). Model comparisons using competitor density instead of biomass as the explanatory variable did not show systematic improvements in model fit (Table S2). Hence, we focus our discussion here on the models using competitor biomass (Fig. 2) and present the results based on competitor density in the Appendix (Fig. S2 and S3).

**Figure 2.**
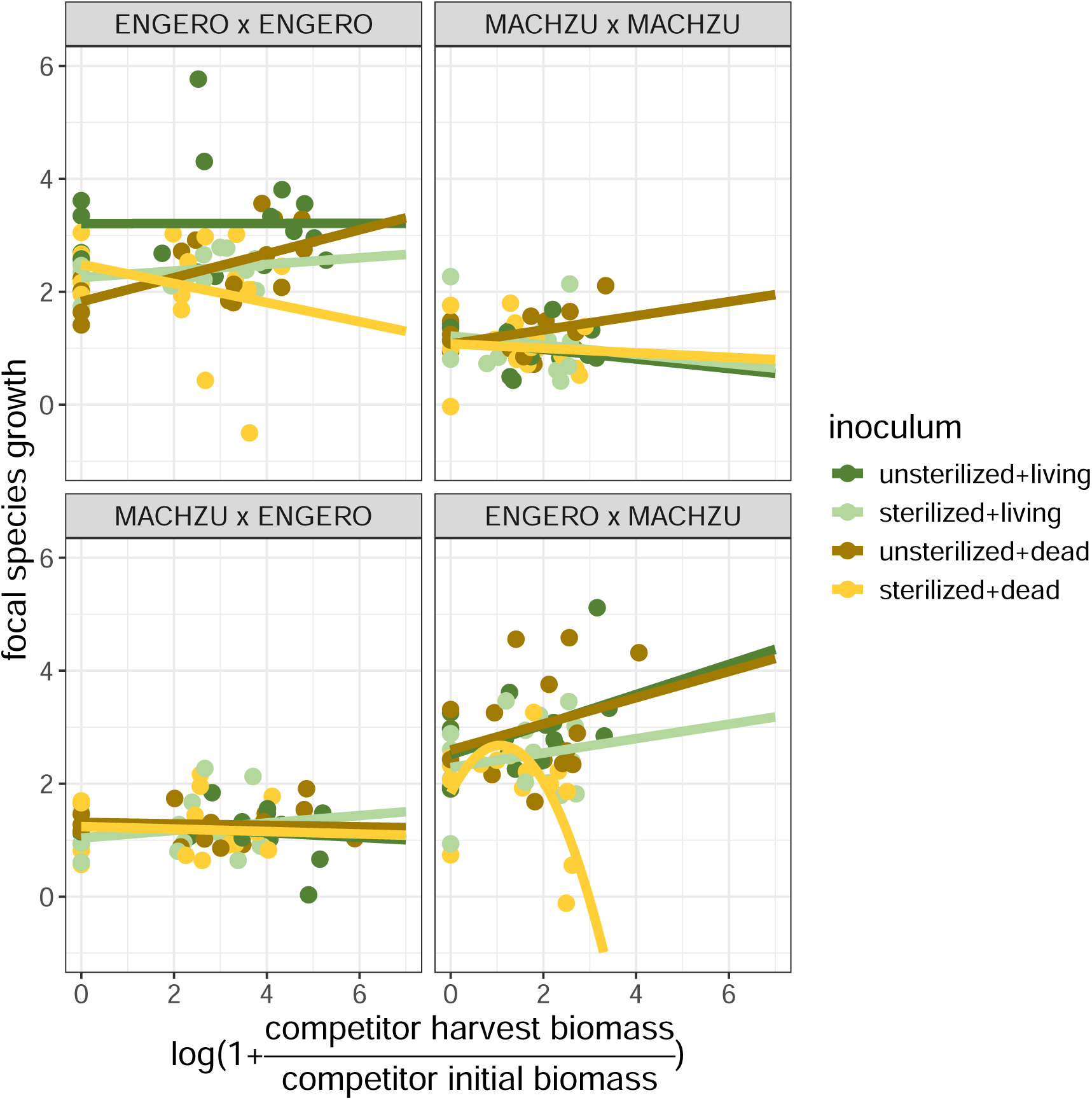
Model fits depicting the effect of plant competitor biomass on the focal individual’s growth. The linear model is fitted for each combination of competing species and inoculum type independently. Panel subtitles indicate the combination of competing species, where the first abbreviation indicates the focal species and the second its competitor (ENGERO = *Engelhardia roxburghiana* and MACHZU = *Machilus zuihoensis*). Inoculum type is a combination of sterilization treatment (unsterilized or sterilized) and host plant status (living or dead), represented by different colors. The y-axis represents the focal individual growth, which is calculated by log-ratio of its dry biomass at harvest to the average initial dry biomass of the species (i.e., log(biomass*_harvest_*/biomass*_initial_*)). The x-axis represents the transformed competitor biomass, which is calculated as the log-transformed harvested dry biomass of the competitors (i.e., log(1 + biomass*_harvest_*/biomass*_initial_*)). Linear or quadratic regression lines are drawn depending on the model selection outcome.

Based on bootstrap samples, we found *E. roxburghiana* generally had greater single individual growth than *M. zuihoensis* across most inocula (comparison between blue and red bars in Fig. 3a). For *M. zuihoensis*, there was no significant difference (i.e., overlapping bootstrap CI) in growth when it grew alone in different inocula. In contrast, *E. roxburghiana* showed a significantly higher single individual growth in unsterilized inocula from living conspecific hosts (dark blue bar; estimate = 3.208, 90% CI: 2.735 – 3.834; Table S3), compared to that in sterilized inocula from living conspecific hosts (estimate = 2.253, 90% CI: 1.196 – 2.461) and in unsterilized inocula from dead conspecific hosts (estimate = 1.826, 90% CI: 1.522 – 2.122).

**Figure 3.**
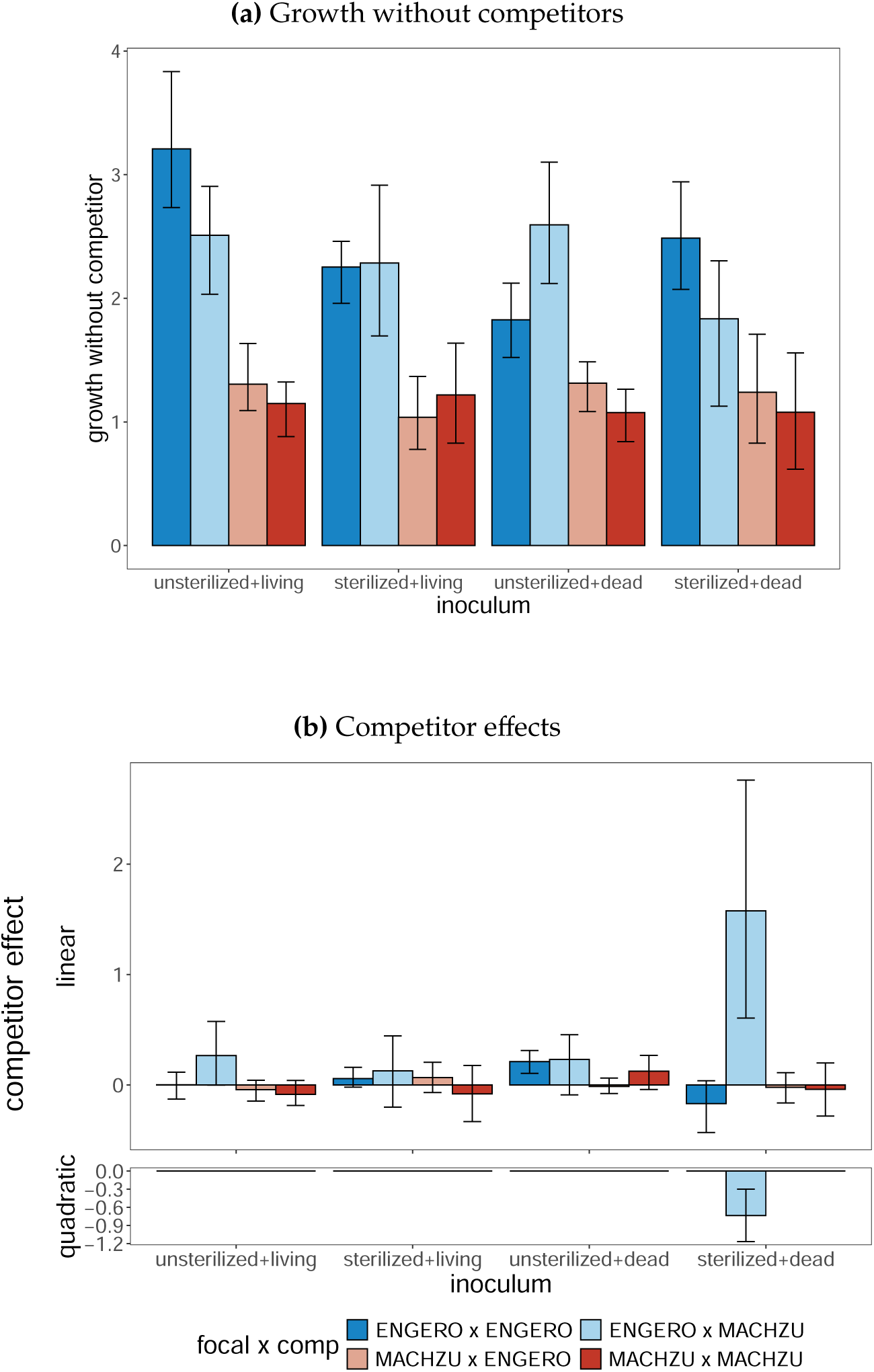
Bootstrap distribution of (a) single-individual growth without competitors and (b) competitor effects for each inoculum type (combination of sterilization treatment and host tree statuses). Bar colors indicate the focal x competitor species combinations (species acronyms: ENGERO = *Engelhardia roxburghiana*; MACHZU = *Machilus zuihoensis*). Bar heights represent (a) the estimated growth without competitor (intercepts) and (b) the estimated effects of competitor biomass on focal species’ growth (linear and quadratic terms), derived from linear regressions of focal species’ growth on transformed competitor biomass (Fig. 2). Error bars represent the 90% bootstrap confidence intervals for each model parameter.

Most of the competitor effects were not significantly different from zero. Two exceptions stood out: (1) the significantly negative quadratic heterospecfic competitor effect on *E. roxburghiana* (light blue bar in Fig. 3b; estimate = −0.732, CI: −1.166 – −0.299; Table S4) in sterilized inocula from dead hosts, and (2) a significantly positive linear conspecific competitor effect among *E. roxburghiana* (dark blue bar; estimate = 0.212, CI: 0.105 – 0.312) in unsterilized inocula from dead hosts, which also differed significantly from the corresponding effect in sterilized inocula (estimate = -0.169, CI: -0.430 – 0.038).

### Robustness of species persistence and competitive outcomes

Invasion analysis based on the selected fitted models revealed the estimated species persistence and competitive outcomes for each inoculum (indicated by solid triangles in Fig. 4a and 4b). Using the bootstrap samples, we assessed the robustness of species persistence and competitive outcomes. Based on the sign of IGRs and our 95% threshold criterion, we found robust inferences of species persistence in some cases. Specifically, for *E. roxburghiana*, 98.9% of bootstrap samples suggested a positive IGR in unsterilized inocula from living hosts, while 99.6% suggested a negative IGR in sterilized inocula from dead hosts (Table 1). For *M. zuihoensis*, no IGRs met the 95% threshold to support a robust inference of persistence. We found notable uncertainty in the inferred competitive outcomes for all inoculum types (Table 2). Nevertheless, we still observed substantial differences in the bootstrap distributions of the inferred species persistence and competitive outcomes among sterilization treatments and host plant statuses.

**Figure 4.**
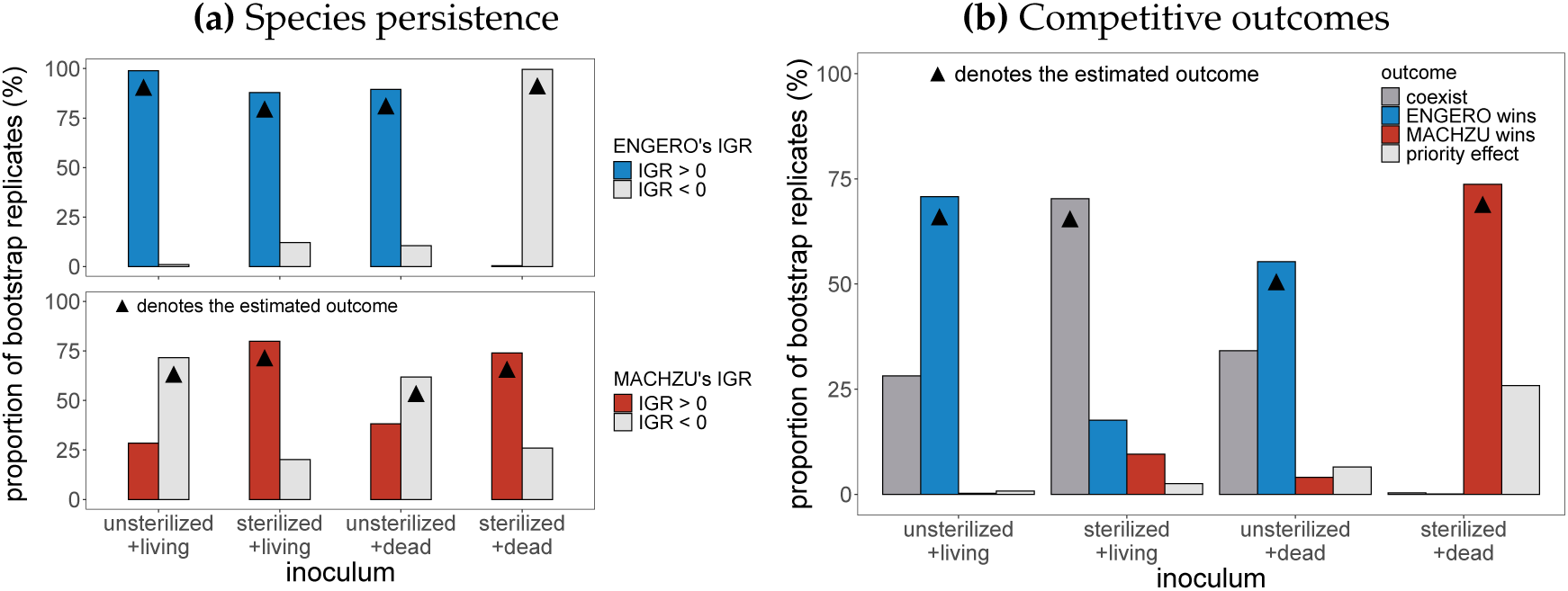
Bootstrap distribution of (a) species persistence and (b) competitive outcome. Bar colors in (a) indicate the sign of the invasion growth rate (IGR) for the two species (blue: IGR > 0 for ENGERO = *Engelhardia roxburghiana*; red: IGR > 0 for MACHZU = *Machilus zuihoensis*; gray: IGR < 0). Bar colors in (b) indicate the competitive outcome (blue: ENGERO wins; red: MACHZU wins; dark gray: coexistence; light grey: priority effect). Bar heights represent the proportion of bootstrap samples suggesting each IGR sign and competitive outcome. Black triangles indicate estimated persistence or coexistence outcomes, i.e., those obtained using the entire dataset as opposed to bootstrap resamples.

**Table 1.** Bootstrap distribution of species persistence associated with each inoculum type (combination of sterilization treatment and host tree statuses). Species acronyms: ENGERO = *Engelhardia roxburghiana*; MACHZU = *Machilus zuihoensis*.

| inoculum | IGR | proportion |  |
| --- | --- | --- | --- |
|  |  | ENGERO | MACHZU |
| unsterilized+living | IGR > 0 | 0.9892 | 0.2845 |
|  | IGR < 0 | 0.0108 | 0.7155 |
| sterilized+living | IGR > 0 | 0.8790 | 0.7984 |
|  | IGR < 0 | 0.1210 | 0.2016 |
| unsterilized+dead | IGR > 0 | 0.8945 | 0.3817 |
|  | IGR < 0 | 0.1055 | 0.6183 |
| sterilized+dead | IGR > 0 | 0.0043 | 0.7404 |
|  | IGR < 0 | 0.9957 | 0.2596 |

**Table 2.** Bootstrap distribution of competitive outcomes associated with each inoculum type (combination of sterilization treatment and host tree status). Species acronyms: ENGERO = *Engelhardia roxburghiana*; MACHZU = *Machilus zuihoensis*.

| inoculum | outcome | n | proportion |
| --- | --- | --- | --- |
| unsterilized+living | ENGERO wins | 7075 | 0.7075 |
|  | MACHZU wins | 28 | 0.0028 |
|  | coexist | 2817 | 0.2817 |
|  | priority effect | 80 | 0.0080 |
| sterilized+living | ENGERO wins | 1761 | 0.1761 |
|  | MACHZU wins | 955 | 0.0955 |
|  | coexist | 7029 | 0.7029 |
|  | priority effect | 255 | 0.0255 |
| unsterilized+dead | ENGERO wins | 5532 | 0.5532 |
|  | MACHZU wins | 404 | 0.0404 |
|  | coexist | 3413 | 0.3413 |
|  | priority effect | 651 | 0.0651 |
| sterilized+dead | ENGERO wins | 8 | 0.0008 |
|  | MACHZU wins | 7369 | 0.7369 |
|  | coexist | 35 | 0.0035 |
|  | priority effect | 2588 | 0.2588 |

By comparing species persistence and competitive outcomes in unsterilized versus sterilized inocula, we were able to detect how microbes mediated seedling competition. Specifically, we found differences in species persistence and competitive outcomes between unsterilized and sterilized inocula collected from either living or dead hosts. For inocula conditioned by living hosts (first and second set of bars in Fig. 4a and 4b), the proportion of bootstrap samples suggesting *M. zuihoensis*’s persistence was lower in unsterilized inocula (i.e., decreased from 79.8% in sterilized inocula to 28.5% in unsterilized inocula; Table 1). This resulted in a lower proportion of coexistence (from 70.3% to 28.2%; Table 2) and a higher proportion of competitive wins by *E. roxburghiana* (from 17.6% to 70.8%) in the unsterilized inocula of living hosts. A similar pattern was observed in inocula collected from dead hosts (third and fourth set of bars in Fig. 4a and 4b). The proportion of bootstrap samples suggesting *M. zuihoensis*’s persistence was also lower in unsterilized inocula (i.e., decreased from 74.0% in sterilized inocula to 38.2% in unsterilized inocula; Table 1). However, unlike its consistently high persistence in inocula collected from living hosts, *E. roxburghiana*’s persistence was different among unsterilized and sterilized inocula collected from dead hosts (0.4% in sterilized inocula but 89.6% in unsterilized inocula). This shift resulted in a lower proportion of competitive wins by *M. zuihoensis* (from 73.7% to 4.0%; Table 2) but more wins by *E. roxburghiana* (from 0.1% to 55.3%) in the unsterilized inocula of dead hosts. These differences between unsterilized and sterilized inocula suggest that soil microbes under both living and dead hosts mediated seedling competition, specifically by lowering the persistence of *M. zuihoensis* and enhancing the success of *E. roxburghiana*.

Furthermore, we determined the temporal change in species persistence and competitive outcomes by comparing treatments with inocula collected from living hosts in 2023 and from hosts that died between 2014 and 2019 (first and third set of bars in Fig. 4a and 4b). First, we found no clear difference in persistence and competitive outcomes between unsterilized inocula collected from living versus dead hosts: *E. roxburghiana* had a consistently higher proportion of persistence (98.9% in living host and 89.5% in dead host inocula; Table 1), while *M. zuihoensis* has a consistently lower proportion of persistence (28.5% in living host and 38.2% in dead host inocula). Accordingly, the unsterilized inocula from different host tree statuses resulted in similar bootstrap distributions of inferred competitive outcomes (Table 2). Digging deeper into the fitted model coefficients, recall that we found *E. roxburghiana* had a greater single individual growth in unsterilized inoculum from living hosts, compared to the one from dead hosts (Fig. 3a; Table S3). Moreover, its conspecific competitor effect shifted from non-significant in inoculum from living hosts (estimate = 0.0008 with 90% CI: −0.127 – 0.116; Fig. 3b; Table S4) to significantly positive in inoculum from dead hosts. Despite these differences, the monoculture equilibrium of *E. roxburghiana* remained similar and its competitive impact on *M. zuihoensis* did not differ significantly across host statuses, resulting in a similar IGR bootstrap distribution for *M. zuihoensis* (see IGR calculation in Data Analysis). Together, these results suggest that microbial mediation of seedling competition remains consistent for 4–9 years after host death, despite detectable differences in single individual growth and competitor effects.

Finally, we found differences in species persistence and competitive outcomes between the sterilized inocula from different host statuses (second and fourth set of bars in Fig. 4a and 4b). Specifically, there was a higher proportion of bootstrap samples suggesting *E. roxburghiana*’s persistence in sterilized inocula from living hosts (87.9%), compared to those from dead hosts (0.4%; Table 1). This difference led to a shift in the bootstrap distribution of competitive outcomes: coexistence in 70.3% and *M. zuihoensis* wins in 9.6% of bootstrap samples in sterilized inocula from living hosts, compared to 0.4% and 73.7%, respectively, in sterilized inocula from dead hosts (Table 2) These results suggest that non-microbial legacies associated with host death can influence seedling competition in sterilized inocula.

## Discussion

By combining field tree mortality data with a density-gradient greenhouse experiment, we revealed the important role of soil microbial communities in mediating seedling competition under both living and dead trees. However, the hypothesis that the effects of PSF on seedling competition would vary through time was not supported by our results. Indeed, we found no differences in microbial-mediated competitive outcomes over 4–9 years, as intact soil microbial communities consistently favored the exclusion of *M. zuihoensis* by *E. roxburghiana*, regardless of whether the inocula were sourced from living or dead trees. These temporally persistent PSF align with Bennett et al. (2022), which showed similar consistent negative effects of soils from dead trees on conspecific seedlings’ survival and growth in a greenhouse experiment, and with Magee et al. (2024), which showed that dead trees impose strong and pervasive negative density dependence on conspecific survival. Notably, a wealth of evidence suggests that key microbial regulators of plant performance, such as oomycete pathogens (Esch and Kobe, 2021), arbuscular mycorrhizal fungi (Varga et al., 2015), and ectomycorrhizal fungi (Shemesh et al., 2023, Mueller et al., 2019), may persist for years to decades after host death through long-lived propagules, providing a mechanistic basis for temporally persistent PSF effects. Adding to these studies, our findings highlight the importance of soil microbial legacies in plant competition in real field systems.

Furthermore, our manipulative experiment suggested that the drivers of PSF may have shifted over time, even when the overall competitive consequences of soil effects were unchanged. In particular, sterilized inoculum treatments allowed us to isolate abiotic effects, i.e., those that did not depend on an active microbial community. In contrast to the unsterilized treatments where it always persisted, *E. roxburghiana* persisted in sterilized inoculum from living but not from dead hosts. We speculate that this shift in competitive outcomes with host plant death was driven primarily by abiotic factors. Despite the low proportion of inoculum (10%) in our soil treatments, plants responded to abiotic differences between the two sterilized inocula, likely reflecting strong nutrient sensitivity in our unfertilized pots. In contrast, our evidence from unsterilized soils suggested that the intact microbial community can mask these underlying abiotic differences, yielding similar competitive outcomes across living versus dead host soils Accordingly, we hypothesize that the temporally persistent PSF observed in our study system, regardless of the temporal change in the sterilized baseline, may reflect the interplay between symbiotic and abiotic factors: selection from a long-lived microbial propagule bank can maintain consistent PSF effects even as other processes, such as nutrient cycling, are altered. This interpretation accords with growing evidence that plants actively select distinct microbial communities (Duhamel et al., 2019) and that such microbial symbioses can maintain plant performance under changing stressors such as nutrient availability (Bogar et al., 2022) and competition (Willing et al., 2024).

Indeed, in line with recent calls (in ’t Zandt et al., 2020) to address soil nutrient availability as a mechanism mediating plant–soil microbe interactions, Van Nuland et al. (2023) combined competition, nitrogen addition, and inoculum sterilization treatments to show how context-dependent interactions between soil nutrients and plant–microbe symbioses jointly determine competitive outcomes between tree species. Our findings add an important temporal dimension to this line of research: persistence of microbial communities after plant death may interact with concurrent changes such as altered nutrient cycling to determine the overall impact of PSFs on seedling competition. Further experiments incorporating microbial community changes after host death (Ou et al., 2024), in tandem with manipulations such as nutrient addition (Van Nuland et al., 2023), nutrient tracing (Bogar et al., 2022), and molecular methods (Janse van Rensburg et al., 2025) will be instrumental in further pinpointing the microbial processes responsible for persistent soil legacies.

Our findings also indicate important species-level differences in PSF and its consequences over time. We always detected greater persistence of *E. roxburghii* in unsterilized soil versus sterilized controls, an effect that was particularly strong in soils from dead hosts. This pattern suggests that *E. roxburghii* benefited from soil microbes, likely due to ectomycorrhizal symbioses providing nutrient benefits (Segnitz et al., 2020) or pathogen protection (Bennett et al., 2017). These persistent positive feedbacks, writ large in the context of a multispecies community (Eppinga et al., 2018), may contribute to the high abundance of *E. roxburghii* at Fushan FDP, as shown for other dominant tree species in species-rich forest ecosystems (Segnitz et al., 2022). On the other hand, our experiment provided stronger support for the persistence of *M. zuihoensis* in sterilized than in unsterilized inocula. This suggests that *M. zuihoensis* experienced negative microbial effects, consistent with previous comparisons showing that arbuscular mycorrhizal plants show more negative PSFs than their ectomycorrhizal competitors (Eagar et al., 2025). Moreover, host death had little impact on bootstrap support for *M. zuihoensis* persistence, regardless of whether the inoculum was sterilized. Nor did we detect any significant changes in the slope or intercept of competition models for *M. zuihoensis* between living versus dead host soils, in contrast to the treatment-specific differences for *E. roxburghii*. We hypothesize that the consistent negative microbial effects experienced by *M. zuihoensis* may reflect its close phylogenetic relationship to the surrounding Lauraceae vegetation at Fushan FDP. In other words, soil conditioning by abundant confamilial neighbors may maintain negative soil microbial effects for *M. zuihoensis* seedlings even after adult hosts die. This parallels results from (Segnitz et al., 2020), who found that arbuscular mycorrhizal trees in a hyperdiverse tropical forest experience negative PSF from close relatives, and emphasizes pathogen spillover from common species as a key mechanism in maintaining microbial effects after plant death. More broadly, our findings point to the importance of further mechanistic work on how PSF persistence varies between host plants, especially in species-rich low-latitude forests where symbiotic traits and phylogenetic relatedness are known to strongly modulate PSFs (Segnitz et al., 2020, Liang et al., 2020).

Using a density gradient, we untangled the nuanced impacts of soil microbes on different aspects of competition. Importantly, complex changes in density dependence were observed for *E. roxburghiana*: we found that soil microbial effects from living trees increased *E. roxburghiana*’s growth alone, while microbial legacies from dead trees resulted in positive (i.e., facilitative) intraspecific density dependence for *E. roxburghiana*. Although these microbial effects influenced *E. roxburghiana* in different manners, both ultimately promoted its dominance during seeding competition when it imposed negative competitor effects on *M. zuihoensis*. Large-scale studies on legacy soil microbial effects have likewise shown changes in density-dependent impacts on demographic rates (e.g., Magee et al. 2024). Thus, we underscore the need to incorporate density-dependent microbial effects, long studied in the context of plant population ecology (Watkinson and Freckleton, 1996), into experimental PSF studies. Meanwhile, model selection identified significant nonlinearity in the response of *E. roxburghiana* competing with *M. zuihoensis* in sterilized soil collected from dead trees (Fig. 3b). Moreover, a significant positive linear effect was observed in *E. roxburghiana*’s conspecific competition in unsterilized dead host soils; as biological realism dictates that performance cannot increase indefinitely with competitor density (Ke and Wan, 2023), this result provides indirect evidence for unobserved nonlinear response. These patterns might indicate the role of microbial communities in shaping the nonlinearity of seedling competition (West, 1996, Willing et al., 2024). As theoretical work has already demonstrated that complex demographic responses may arise from the underlying biology of species interactions (e.g., Holland and DeAngelis 2010, Letten and Stouffer 2019), studying these nonlinearities is especially important in light of the long-standing debate regarding appropriate functional forms for competitive responses (Ayala et al., 1973) and their coexistence implications (Armitage, 2024, Hatton et al., 2024). We emphasize the utility of density gradient experiments in providing a more complete picture to link PSF mechanisms to realistic, potentially nonlinear, competition functional responses.

Nonetheless, our findings also highlight methodological challenges in quantifying density-dependent microbial effects. Similar to previous studies quantifying plant coexistence (e.g., Chu and Adler 2015), including those from the perspective of plant–soil interactions (Willing et al., 2024), we found that estimated competitive effects were close to zero in many cases, contributing to considerable uncertainty in the predicted competitive outcome. This likely reflects the difficulty of choosing density treatments that allow accurate estimation of monoculture equilibrium (Ke and Wan, 2023). In other cases, challenges in model-fitting may be related to potential nonlinearities in competitive responses. For instance, though one nonlinear term was retained and bootstrapping indicated robust support for some positive competitive responses (Fig. 3b), parameter uncertainty was relatively large. While we were still able to apply our invasion analysis-based approach to qualitatively predict coexistence, our results emphasize the importance of assessing model robustness. Accordingly, we echo recent recommendations to assess uncertainty via model selection (Armitage, 2024) and bootstrapping (as implemented here; Fig. 4), or other techniques including error propagation (Yan et al., 2022) or Bayesian statistics (Terry and Armitage, 2024).

In summary, by combining long-term field data with a density-gradient greenhouse experiment, we showed that microbial effects can persist for 4–9 years after host death and consistently affect seedling competition. Our results also highlight the potential for interactions between consistent microbial communities and temporal shifts in abiotic conditions. Moreover, we revealed the role of soil microbes in mediating the nonlinear density dependence in seedling competitive responses. Crucially, these microbial effects act through density-dependent competition functions, which highlights that assessing plant–soil interactions using only single individuals may overlook important features of PSF. Looking forward, we recommend combining density gradient experiments with robust methods for model selection and uncertainty estimation. Moreover, future work may gain further mechanistic insight by pairing density dependent plant–soil feedback experiments with direct measurements of microbial turnover. Together, these approaches will improve our ability to predict plant community dynamics through the lens of plant–soil microbe interactions.

## Acknowledgements

This experiment would not have been possible without the support of many people. We thank Yi Sun, Shu-Ping Wang, Hsun-Hung Chu, Yu-Pei Tseng, Chin-Te Tsai, Hsiang-Chih Lo, Sheng-Yueh Liang, Ni-Chen Lin, and others who assisted with soil collection, planting, and harvest. We thank members of the Ke Lab for providing valuable feedback on the early draft of the manuscript. The Taiwan Forestry and Nature Conservation Agency, the Taiwan Forestry Research Institute, and the Ministry of Science and Technology of Taiwan support the Fushan Forest Dynamics Plot (FDP). This study is supported by the Yushan Fellow Program, Ministry of Education, Taiwan (MOE-110-YSFAG-0003-001-P1), the National Science and Technology Council, Taiwan (NSTC 113-2811-B-002-118 and NSTC 114-2628-B-002-023-), and National Taiwan University (Career Development Projects NTU-114L7867 and postdoctoral grant 112L4000-1).

## Statements and Declarations

### Author Contribution

PJK and JW conceived the study; CLH conducted the study and analyzed the data with assistance from JW and SW; CLH wrote the first draft of the manuscript with help from JW and PJK; CCY provided tree mortality survey data in Fushan FDP; all authors contributed to the final version of the manuscript.

### Data Availability

Should the manuscript be accepted, all data and computer scripts supporting the results will be archived in an appropriate public repository with the DOI included at the end of the article.

---

**Figure S1.**
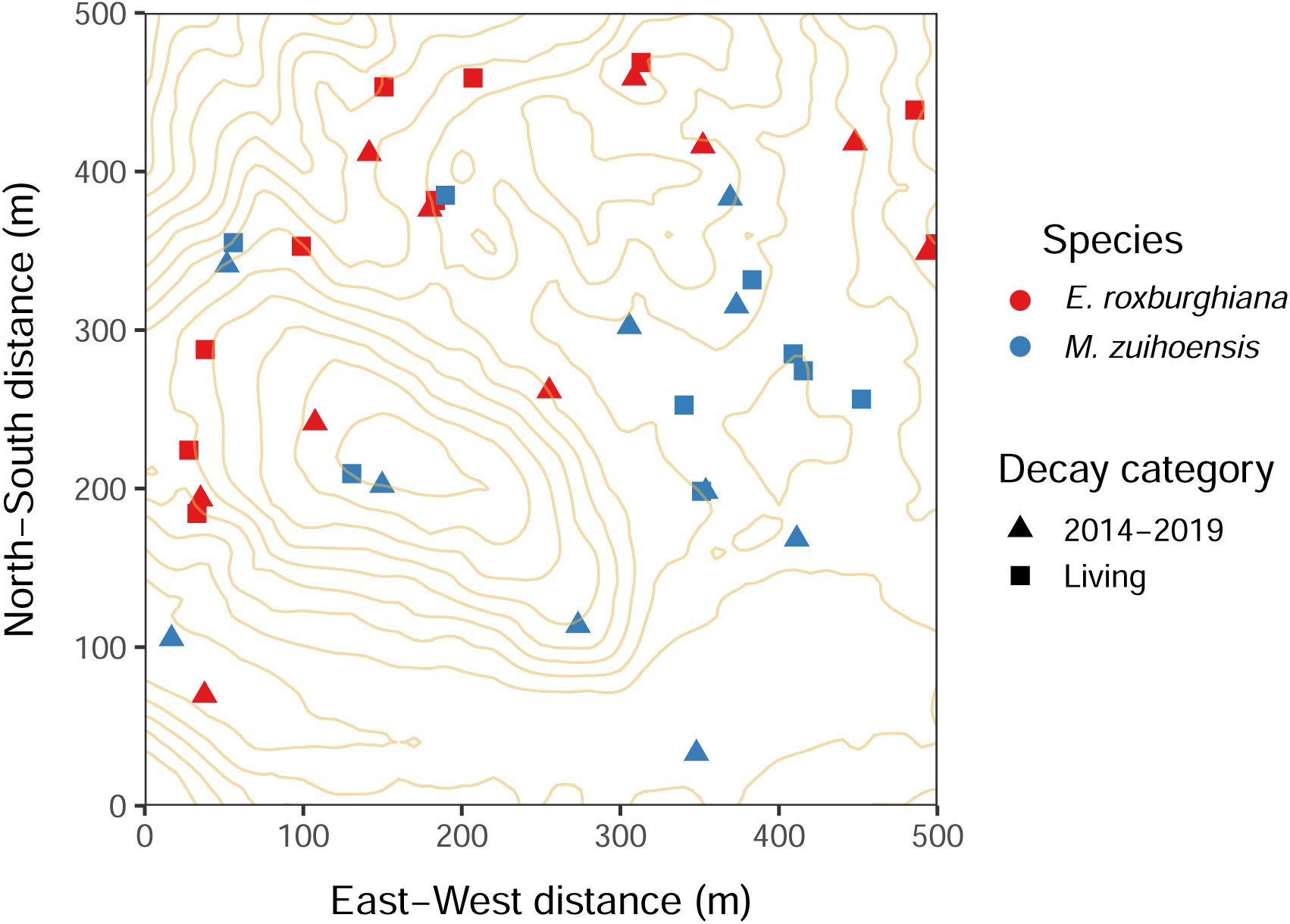
Sampling map in Fushan Forest Dynamics Plot. Blue points represent *Engelhardia roxburghiana* and red points represent *Machilus zuihoensis*. Triangular shapes represent individuals recorded as dead during the 2014–2019 period, while squared shapes represent individuals recorded as living during our soil sampling in 2023.

**Figure S2.**
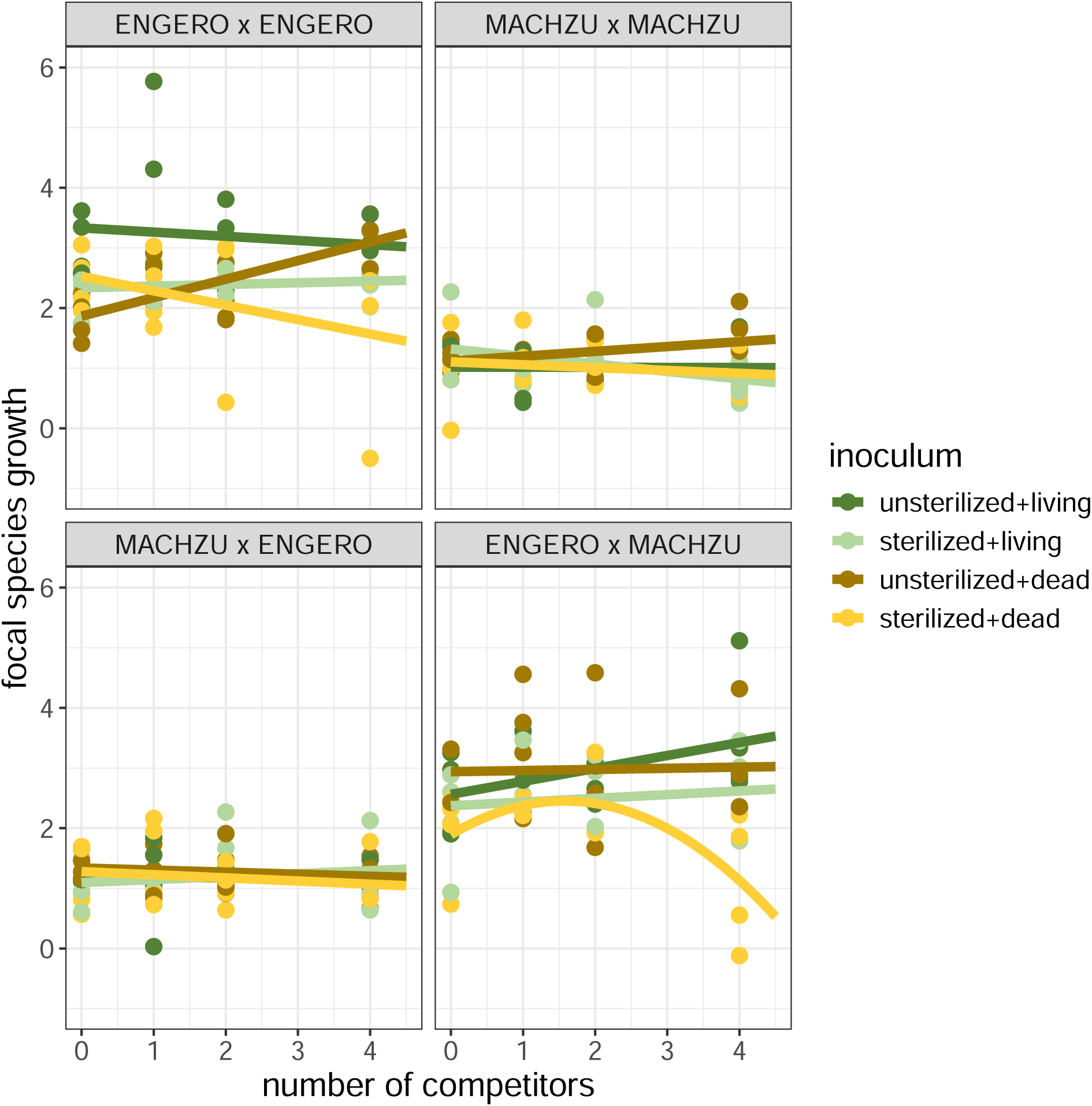
Model fit of plant competitor density on focal individual’s growth. The linear model is fitted for each combination of competing species and inoculum type independently. Each panel has a subtitle on the top, indicating a combination of competing species, with the first one as the focal species and the second one as its competitor (ENGERO = *Engelhardia roxburghiana* and MACHZU = *Machilus zuihoensis*). Inoculum type is a combination of sterilization treatment (unsterilized or sterilized) and host plant status (living or dead), represented by different colors. The y-axis represents the focal individual growth, which is calculated by log-ratio of its dry biomass at harvest to the average initial dry biomass of the species (i.e., log(biomass*_harvest_*/biomass*_initial_*)). The x-axis represents the number of competitors. Linear or quadratic regression lines are drawn based on the model selection results.

**Figure S3.**
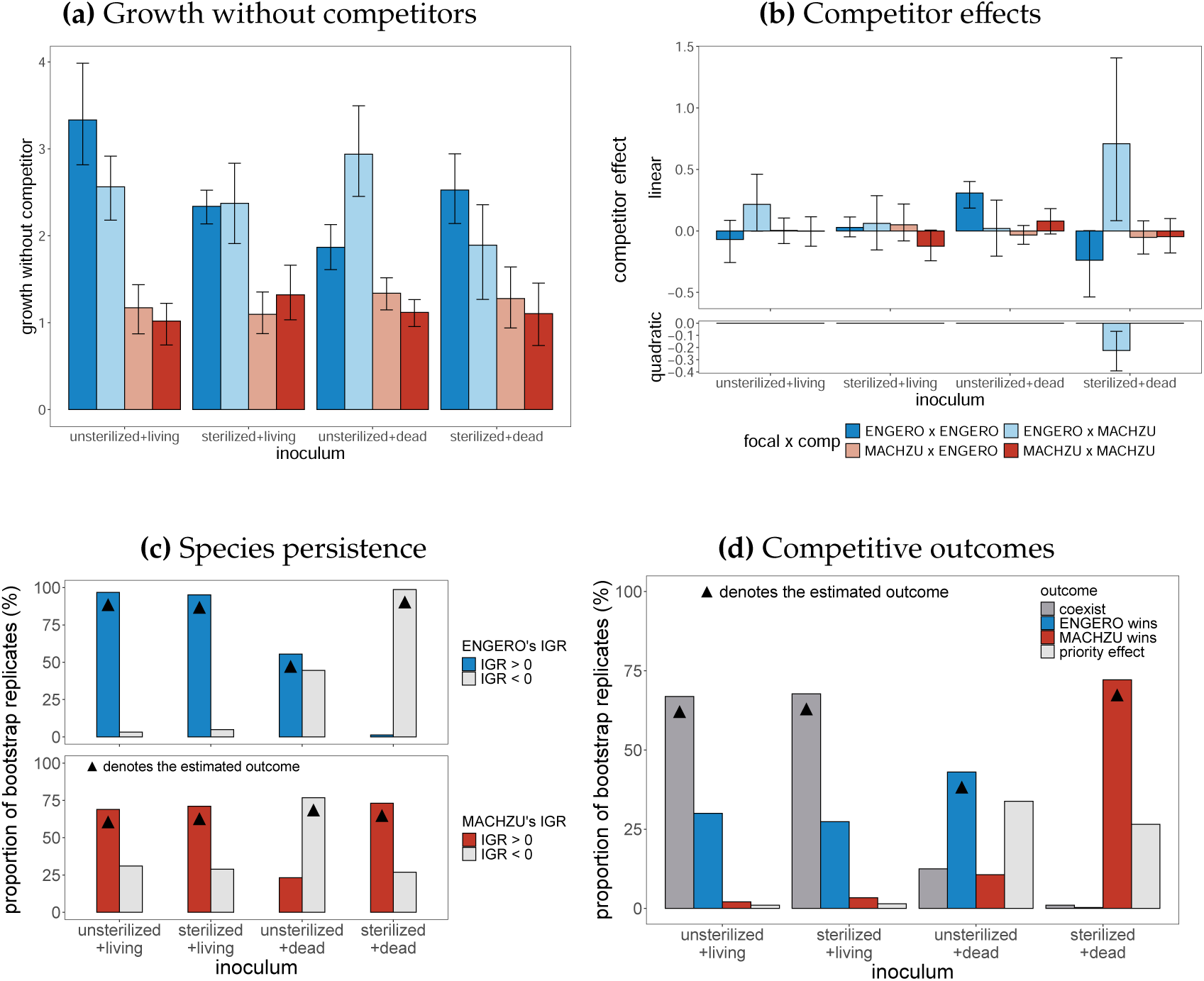
Bootstrap distribution of single individual growth, competitor effects, species persistence and competitive outcomes when regressing focal species growth on the number of competitors. See more descriptions of the diagrams in Fig. 3 and 4.

**Table S1.** Results of F-test of significance of linear and quadratic plant-plant interactions when regressing focal species growth on transformed competitor biomass. Inocula are combination of sterilization treatment and host tree status. Species acronyms: ENGERO = *Engelhardia roxburghiana*; MACHZU = *Machilus zuihoensis*.

| inoculum | focal x comp | Res.Df | RSS | Df | Sum.of.Sq | F | P value |
| --- | --- | --- | --- | --- | --- | --- | --- |
| unsterilized+living | ENGERO x ENGERO | 15 | 12.04723 | NA | NA | NA | NA |
|  |  | 14 | 12.04720 | 1 | 0.00004 | 0.00005 | 0.99474 |
|  |  | 13 | 11.39456 | 1 | 0.65263 | 0.74458 | 0.40383 |
|  | ENGERO x MACHZU | 15 | 7.84058 | NA | NA | NA | NA |
|  |  | 14 | 6.34676 | 1 | 1.49382 | 3.29714 | 0.09253 |
|  |  | 13 | 5.88986 | 1 | 0.45690 | 1.00847 | 0.33360 |
|  | MACHZU x ENGERO | 15 | 2.72337 | NA | NA | NA | NA |
|  |  | 14 | 2.61784 | 1 | 0.10553 | 0.61039 | 0.44864 |
|  |  | 13 | 2.24756 | 1 | 0.37029 | 2.14175 | 0.16709 |
|  | MACHZU x MACHZU | 15 | 1.91105 | NA | NA | NA | NA |
|  |  | 14 | 1.77775 | 1 | 0.13330 | 1.12678 | 0.30779 |
|  |  | 13 | 1.53796 | 1 | 0.23979 | 2.02685 | 0.17810 |
| sterilized+living | ENGERO x ENGERO | 15 | 1.16474 | NA | NA | NA | NA |
|  |  | 14 | 1.05400 | 1 | 0.11074 | 1.36627 | 0.26344 |
|  |  | 13 | 1.05367 | 1 | 0.00034 | 0.00416 | 0.94953 |
|  | ENGERO x MACHZU | 15 | 7.02878 | NA | NA | NA | NA |
|  |  | 14 | 6.77139 | 1 | 0.25740 | 0.52823 | 0.48024 |
|  |  | 13 | 6.33467 | 1 | 0.43672 | 0.89623 | 0.36106 |
|  | MACHZU x ENGERO | 15 | 3.52007 | NA | NA | NA | NA |
|  |  | 14 | 3.38604 | 1 | 0.13403 | 0.53025 | 0.47942 |
|  |  | 13 | 3.28600 | 1 | 0.10004 | 0.39577 | 0.54018 |
|  | MACHZU x MACHZU | 15 | 3.67568 | NA | NA | NA | NA |
|  |  | 14 | 3.57605 | 1 | 0.09964 | 0.38408 | 0.54614 |
|  |  | 13 | 3.37244 | 1 | 0.20361 | 0.78486 | 0.39175 |
| unsterilized+dead | ENGERO x ENGERO | 15 | 6.14248 | NA | NA | NA | NA |
|  |  | 14 | 4.01687 | 1 | 2.12561 | 6.95227 | 0.02052 |
|  |  | 13 | 3.97467 | 1 | 0.04220 | 0.13801 | 0.71625 |
|  | ENGERO x MACHZU | 15 | 12.67356 | NA | NA | NA | NA |
|  |  | 14 | 11.46389 | 1 | 1.20967 | 1.43611 | 0.25216 |
|  |  | 13 | 10.95016 | 1 | 0.51374 | 0.60991 | 0.44881 |
|  | MACHZU x ENGERO | 15 | 1.64174 | NA | NA | NA | NA |
|  |  | 14 | 1.63249 | 1 | 0.00925 | 0.07842 | 0.78386 |
|  |  | 13 | 1.53360 | 1 | 0.09889 | 0.83825 | 0.37657 |
|  | MACHZU x MACHZU | 15 | 1.79523 | NA | NA | NA | NA |
|  |  | 14 | 1.52957 | 1 | 0.26565 | 3.07604 | 0.10298 |
|  |  | 13 | 1.12271 | 1 | 0.40687 | 4.71120 | **0.04908** |
| sterilized+dead | ENGERO x ENGERO | 15 | 13.79237 | NA | NA | NA | NA |
|  |  | 14 | 12.84345 | 1 | 0.94893 | 0.96058 | 0.34493 |
|  |  | 13 | 12.84234 | 1 | 0.00111 | 0.00112 | 0.97383 |
|  | ENGERO x MACHZU | 15 | 10.95750 | NA | NA | NA | NA |
|  |  | 14 | 10.23641 | 1 | 0.72108 | 1.49897 | 0.24256 |
|  |  | 13 | 6.25369 | 1 | 3.98273 | 8.27918 | **0.01296** |
|  | MACHZU x ENGERO | 15 | 3.94046 | NA | NA | NA | NA |
|  |  | 14 | 3.92284 | 1 | 0.01762 | 0.05980 | 0.81063 |
|  |  | 13 | 3.83112 | 1 | 0.09172 | 0.31122 | 0.58641 |
|  | MACHZU x MACHZU | 15 | 3.18043 | NA | NA | NA | NA |
|  |  | 14 | 3.15597 | 1 | 0.02445 | 0.10642 | 0.74944 |
|  |  | 13 | 2.98713 | 1 | 0.16885 | 0.73482 | 0.40685 |

**Table S2.** AIC and BIC values for model selection. The numbers (1 or 2) that follow AIC and BIC in the column represent the model type, with 1 representing the model that only considers the intercept and the linear term, and 2 representing the model that considers the quadratic term. The column’s subtitle indicates that competitor biomass or density is used in regressing focal species growth. The inoculum column represents the combination of sterilization treatment and host tree status. Species acronyms: ENGERO = *Engelhardia roxburghiana*; MACHZU = *Machilus zuihoensis*.

| inoculum | focal x comp | biomass |  |  |  | density |  |  |  |
| --- | --- | --- | --- | --- | --- | --- | --- | --- | --- |
|  |  | AIC.1 | AIC.2 | BIC.1 | BIC.2 | AIC.1 | AIC.2 | BIC.1 | BIC.2 |
| unsterilized<br>+living | ENGERO x ENGERO | **46.866** | 47.975 | **49.184** | 51.065 | 46.638 | 48.456 | 48.956 | 51.547 |
|  | ENGERO x MACHZU | **36.612** | 37.416 | **38.929** | 40.507 | 36.281 | 37.831 | 38.599 | 40.921 |
|  | MACHZU x ENGERO | 22.442 | **22.002** | **24.760** | 25.092 | 23.073 | 24.991 | 25.390 | 28.081 |
|  | MACHZU x MACHZU | 16.250 | **15.932** | **18.568** | 19.022 | 17.406 | 13.803 | 19.724 | 16.894 |
| sterilized<br>+living | ENGERO x ENGERO | **7.886** | 9.881 | **10.204** | 12.971 | 9.117 | 9.691 | 11.435 | 12.782 |
|  | ENGERO x MACHZU | **37.648** | 38.581 | **39.966** | 41.672 | 37.940 | 39.044 | 40.258 | 42.135 |
|  | MACHZU x ENGERO | **26.559** | 28.079 | **28.877** | 31.170 | 26.788 | 26.885 | 29.106 | 29.975 |
|  | MACHZU x MACHZU | **27.433** | 28.495 | **29.750** | 31.585 | 25.309 | 26.620 | 27.627 | 29.710 |
| unsterilized<br>+dead | ENGERO x ENGERO | **29.293** | 31.124 | **31.610** | 34.214 | 23.733 | 25.189 | 26.051 | 28.279 |
|  | ENGERO x MACHZU | **46.072** | 47.338 | **48.390** | 50.429 | 47.661 | 49.418 | 49.979 | 52.509 |
|  | MACHZU x ENGERO | **14.886** | 15.887 | **17.204** | 18.977 | 14.572 | 16.554 | 16.890 | 19.644 |
|  | MACHZU x MACHZU | 13.844 | **10.896** | 16.162 | **13.987** | 14.253 | 15.114 | 16.570 | 18.205 |
| sterilized<br>+dead | ENGERO x ENGERO | **47.890** | 49.889 | **50.208** | 52.979 | 46.520 | 48.408 | 48.837 | 51.499 |
|  | ENGERO x MACHZU | 44.260 | **38.375** | 46.578 | **41.466** | 42.591 | 39.424 | 44.909 | 42.514 |
|  | MACHZU x ENGERO | **28.914** | 30.535 | **31.231** | 33.625 | 28.595 | 30.583 | 30.913 | 33.673 |
|  | MACHZU x MACHZU | **25.433** | 26.554 | **27.751** | 29.644 | 25.155 | 26.460 | 27.473 | 29.550 |

**Table S3.**
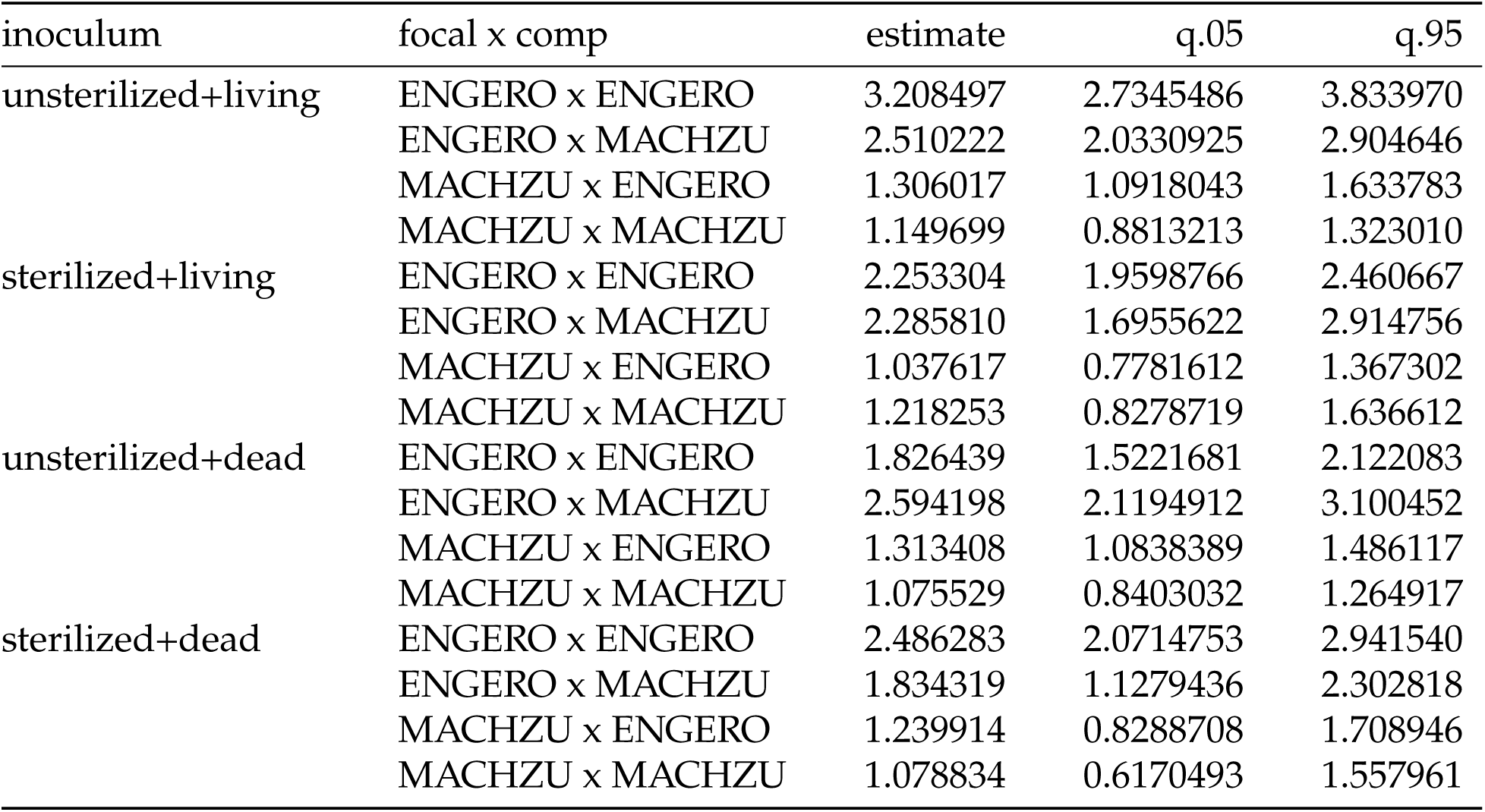
Bootstrap distribution of single individual growth associated with each inoculum (combination of sterilization treatment and host tree status) and focal-competitor species combination (ENGERO = *Engelhardia roxburghiana*; MACHZU = *Machilus zuihoensis*). This table corresponds to Fig. 3a.

| inoculum | focal x comp | estimate | q.05 | q.95 |
| --- | --- | --- | --- | --- |
| unsterilized+living | ENGERO x ENGERO | 3.208497 | 2.7345486 | 3.833970 |
|  | ENGERO x MACHZU | 2.510222 | 2.0330925 | 2.904646 |
|  | MACHZU x ENGERO | 1.306017 | 1.0918043 | 1.633783 |
|  | MACHZU x MACHZU | 1.149699 | 0.8813213 | 1.323010 |
| sterilized+living | ENGERO x ENGERO | 2.253304 | 1.9598766 | 2.460667 |
|  | ENGERO x MACHZU | 2.285810 | 1.6955622 | 2.914756 |
|  | MACHZU x ENGERO | 1.037617 | 0.7781612 | 1.367302 |
|  | MACHZU x MACHZU | 1.218253 | 0.8278719 | 1.636612 |
| unsterilized+dead | ENGERO x ENGERO | 1.826439 | 1.5221681 | 2.122083 |
|  | ENGERO x MACHZU | 2.594198 | 2.1194912 | 3.100452 |
|  | MACHZU x ENGERO | 1.313408 | 1.0838389 | 1.486117 |
|  | MACHZU x MACHZU | 1.075529 | 0.8403032 | 1.264917 |
| sterilized+dead | ENGERO x ENGERO | 2.486283 | 2.0714753 | 2.941540 |
|  | ENGERO x MACHZU | 1.834319 | 1.1279436 | 2.302818 |
|  | MACHZU x ENGERO | 1.239914 | 0.8288708 | 1.708946 |
|  | MACHZU x MACHZU | 1.078834 | 0.6170493 | 1.557961 |

**Table S4.** Bootstrap distribution of competitor effects associated with each inoculum (combination of sterilization treatment and host tree status) and focal-competitor species combination (ENGERO = *Engelhardia roxburghiana*; MACHZU = *Machilus zuihoensis*). This table corresponds to Fig. 3b

| inoculum | focal x comp | estimate | q.05 | q.95 |
| --- | --- | --- | --- | --- |
| 1st order |  |  |  |  |
| unsterilized+living | ENGERO x ENGERO | 0.00084 | -0.12709 | 0.11628 |
|  | ENGERO x MACHZU | 0.26702 | -0.00222 | 0.57593 |
|  | MACHZU x ENGERO | -0.04299 | -0.14597 | 0.04289 |
|  | MACHZU x MACHZU | -0.08473 | -0.18605 | 0.04061 |
| sterilized+living | ENGERO x ENGERO | 0.05824 | -0.01985 | 0.15934 |
|  | ENGERO x MACHZU | 0.12783 | -0.20064 | 0.44480 |
|  | MACHZU x ENGERO | 0.06629 | -0.06817 | 0.20471 |
|  | MACHZU x MACHZU | -0.08125 | -0.33155 | 0.17669 |
| unsterilized+dead | ENGERO x ENGERO | 0.21198 | 0.10514 | 0.31236 |
|  | ENGERO x MACHZU | 0.23200 | -0.08931 | 0.45480 |
|  | MACHZU x ENGERO | -0.01303 | -0.07826 | 0.06195 |
|  | MACHZU x MACHZU | 0.12498 | -0.04075 | 0.26733 |
| sterilized+dead | ENGERO x ENGERO | -0.16916 | -0.43025 | 0.03825 |
|  | ENGERO x MACHZU | 1.57592 | 0.60628 | 2.76059 |
|  | MACHZU x ENGERO | -0.02184 | -0.16245 | 0.11063 |
|  | MACHZU x MACHZU | -0.03978 | -0.28183 | 0.19912 |
| 2nd order |  |  |  |  |
| sterilized+dead | ENGERO x MACHZU | -0.73233 | -1.16566 | -0.29922 |

**Table S5.** Bootstrap distribution of species persistence when regressing focal species growth on number of competitors. The inoculum column represents combinations of sterilization treatment and host tree status. Species acronyms: ENGERO = *Engelhardia roxburghiana*; MACHZU = *Machilus zuihoensis*. IGR = invasion growth rate.

| inoculum | IGR | proportion |  |
| --- | --- | --- | --- |
|  |  | ENGERO | MACHZU |
| unsterilized+living | IGR > 0 | 0.9713 | 0.6967 |
|  | IGR < 0 | 0.0287 | 0.3033 |
| sterilized+living | IGR > 0 | 0.9563 | 0.7111 |
|  | IGR < 0 | 0.0437 | 0.2889 |
| unsterilized+dead | IGR > 0 | 0.5600 | 0.2229 |
|  | IGR < 0 | 0.4400 | 0.7771 |
| sterilized+dead | IGR > 0 | 0.0145 | 0.7263 |
|  | IGR < 0 | 0.9855 | 0.2737 |

**Table S6.**
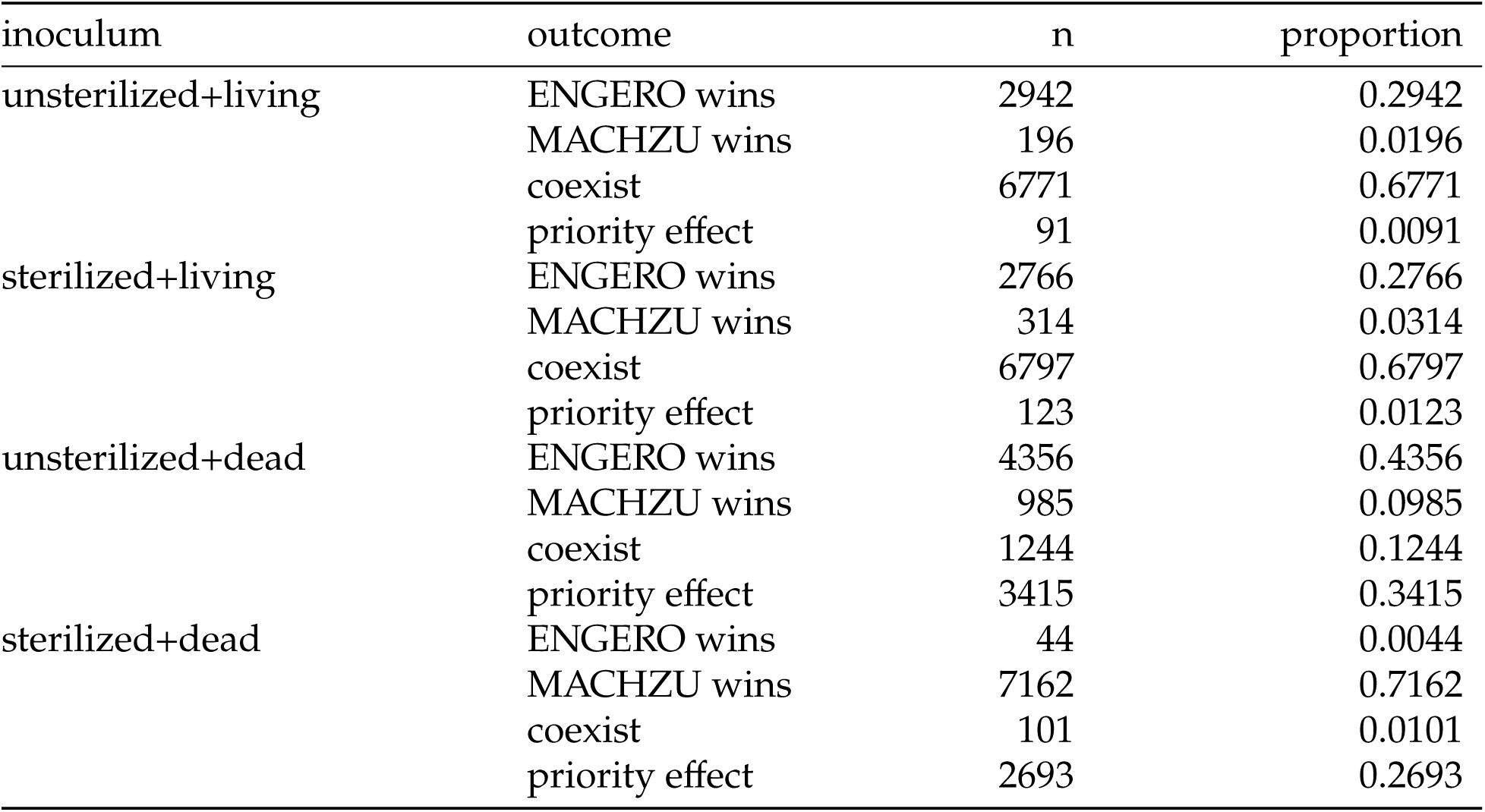
Bootstrap distribution of competitive outcome when regressing focal species growth on the number of competitors. The inoculum column represents combinations of sterilization treatment and host tree status. Species acronyms: ENGERO = *Engelhardia roxburghiana*; MACHZU = *Machilus zuihoensis*.

| inoculum | outcome | n | proportion |
| --- | --- | --- | --- |
| unsterilized+living | ENGERO wins | 2942 | 0.2942 |
|  | MACHZU wins | 196 | 0.0196 |
|  | coexist | 6771 | 0.6771 |
|  | priority effect | 91 | 0.0091 |
| sterilized+living | ENGERO wins | 2766 | 0.2766 |
|  | MACHZU wins | 314 | 0.0314 |
|  | coexist | 6797 | 0.6797 |
|  | priority effect | 123 | 0.0123 |
| unsterilized+dead | ENGERO wins | 4356 | 0.4356 |
|  | MACHZU wins | 985 | 0.0985 |
|  | coexist | 1244 | 0.1244 |
|  | priority effect | 3415 | 0.3415 |
| sterilized+dead | ENGERO wins | 44 | 0.0044 |
|  | MACHZU wins | 7162 | 0.7162 |
|  | coexist | 101 | 0.0101 |
|  | priority effect | 2693 | 0.2693 |

